# Regulation of caveolae through cholesterol-depletion dependent tubulation by PACSIN2/Syndapin II

**DOI:** 10.1101/2020.03.25.008854

**Authors:** Aini Gusmira Amir, Kazuhiro Takemura, Kyoko Hanawa-Suetsugu, Kayoko Oono-Yakura, Kazuma Yasuhara, Akio Kitao, Shiro Suetsugu

**Affiliations:** Division of Biological Science, Nara Institute of Science and Technology; Tokyo Institute of Technology; Division of Material Science, Nara Institute of Science and Technology

## Abstract

The membrane shaping ability of PACSIN2 via its FCH-BAR (F-BAR) domain has been shown to be essential for caveolar morphogenesis, presumably through the shaping of the caveolar neck. Caveolar membrane contains abundant levels of cholesterol. However, the role of cholesterol in PACSIN2-mediated membrane deformation remains unclear. We show that the binding of PACSIN2 to the membrane could be negatively regulated by the amount of cholesterol in the membrane. We prepared a reconstituted membrane based on the lipid composition of caveolae. The reconstituted membrane with cholesterol had a weaker affinity to the F-BAR domain of PACSIN2 than the membrane without cholesterol, presumably due to a decrease in electrostatic charge density. Consistently, the acute depletion of cholesterol from the plasma membrane resulted in the appearance of PACSIN2-localized tubules with caveolin-1 at their tips, suggesting that the presence of cholesterol inhibited the prominent membrane tubulation by PACSIN2. The tubules induced by PACSIN2 were suggested to be an intermediate of caveolae endocytosis. Consistently, the removal of caveolae from the plasma membrane upon cholesterol depletion was diminished in the cells deficient in PACSIN2. These data suggested that PACSIN2 mediated the caveolae internalization dependently on the amount of cholesterol at the plasma membrane, providing a possible mechanism for the cholesterol-dependent regulation of caveolae.

## INTRODUCTION

The Bin-Amphiphysin-Rvs (BAR) domain superfamily of proteins has been shown to play major roles in the shaping of the cell membranes (Daumke et al., 2014; Doherty and McMahon, 2009; Nishimura et al., 2018; Simunovic et al., 2015; Suetsugu et al., 2014). Fes/CIP4 homology (FCH)-BAR (F-BAR) domain proteins, PACSINs (also known as syndapins), are localized to invaginations, such as endocytic sites, including caveolae (Hansen et al., 2011; Koch et al., 2012; Senju et al., 2011). The structural analysis of PACSIN2 F-BAR domains revealed that the F-BAR domains had a positively charged concave surface for membrane invagination, analogous to other BAR and F-BAR domains, that binds to negatively charged lipids including phosphatidylserine (PS) via electrostatic interaction (Rao et al., 2010; Shimada et al., 2010). The membrane binding of PACSIN2 did not appear to be specifically dependent on particular negatively-charged lipids, such as phosphatidylinositol 4-phosphate (PI(4)P) and phosphatidylinositol 4,5-bisphosphate (PI(4,5)P_2_) (Dharmalingam et al., 2009). The F-BAR domains of PACSINs have specific hydrophobic loops that protrude on the concave membrane-binding surface of the structure, which are inserted into the hydrophobic region of the membrane (Shimada et al., 2010).

The membrane of caveolae is composed of several types of lipids, which are cholesterol and phospholipids (Hubert et al., 2020; Moren et al., 2012; Murata et al., 1995; Ortegren et al., 2004; Schlegel et al., 1999). Cholesterol is highly enriched in caveolae (Ortegren et al., 2004; Razani et al., 2002; Smart et al., 1999). Up to 41% of membrane lipids of caveolae in adipocytes reported as cholesterol, which is higher than the typical amount of cholesterol outside caveolae (approximately 22%) (Ortegren et al., 2004). Furthermore, cholesterol is an essential component of caveolae, because cholesterol depletion impairs the morphology of caveolae (Breen et al., 2012; Dreja et al., 2002; Murata et al., 1995; Parpal et al., 2001; Razani et al., 2002; Smart et al., 1999). Due to cholesterol enrichment, caveolae are sometimes considered to be subsets of lipid rafts with caveolin protein localization (Patel and Insel, 2009; Pike, 2003; Razani et al., 2002).

Phospholipids are composed of two fatty acids and one hydrophilic group, such as serine, ethanolamine, choline, and inositol. Phospholipids are primarily classified by their hydrophilic group. The major phospholipids that compose the cellular membrane are phosphatidylethanolamine (PE), phosphatidylcholine (PC), phosphatidylserine (PS), and phosphatidylinositol (PI). PC is widely known to be the most abundant phospholipid in the cell membrane, comprising 41-57 mol % of total glycerophospholipids (van Meer et al., 2008; Yang et al., 2018). PC is also a prominent phospholipid in caveolae (Huot et al., 2010; Pike et al., 2005; Smart et al., 1999), while PS has been found to be abundant in the cytoplasmic leaflet of caveolae (Fairn et al., 2011; Pike et al., 2005). The depletion of PS was shown to result in the loss of caveolar morphology (Hirama et al., 2017). The main fatty acids of caveolar phospholipids are oleic acid (C18:1), palmitic acid (C16:0) (Cai et al., 2013; Huot et al., 2010), and stearic acid (C18:0) (Cai et al., 2013). The 1-palmitoyl-2-oleoyl (16:0-18:1) PC (POPC) and the 1-palmitoyl-2-oleoyl (16:0-18:1) PS (POPS) are abundant phosphatidylcholine and phosphatidylserine in the caveolar membrane, respectively (Pike et al., 2005).

The structural proteins of caveolae are considered to be caveolins (Rothberg et al., 1992), which interact with cholesterol (Murata et al., 1995) and cavins (Hill et al., 2008). Furthermore, palmitic acid and stearic acid are the predominant fatty acids that bind to caveolin-1 (Cai et al., 2013). A caveola contains approximately 150 caveolin proteins (Khater et al., 2019; Pelkmans and Zerial, 2005; Tachikawa et al., 2017), providing a possible explanation for the enrichment of cholesterol in caveolae. PACSIN2 is localized to the neck of caveolae and has been suggested to interact with caveolin-1 through their F-BAR domain (Senju et al., 2011). Approximately 35-48% of endogenous caveolin-1 has been found to colocalize with PACSIN2, implying that PACSIN2 is involved in a subset of caveolae (Hansen et al., 2011; Senju et al., 2011). The knockdown of PACSIN2 in HeLa cells was found to impair the morphology of caveolin-1-associated membranes (Hansen et al., 2011; Senju et al., 2011; Senju et al., 2015). The tension applied to the plasma membrane induces the flattening of caveolae (Sinha et al., 2011). Upon the application of tension, PACSIN2 is removed from caveolae by the protein kinase C (PKC)-mediated phosphorylation of PACSIN2 (Senju et al., 2015). The PKC-phosphorylated PACSIN2 has a weaker affinity to liposomes made of the total lipid fraction of bovine brain, which has been widely used for the study of BAR domains (Senju et al., 2015; Senju and Suetsugu, 2015).

The above mentioned studies showed that PACSIN2 binds to the membrane via its F-BAR domains. However, the lipid characteristics of the caveolae have not yet been studied for membrane deformation by PACSIN2. Furthermore, the effect of cholesterol has not been specifically studied in the model membrane used in *in vitro* assays, nor has tension been applied to the membranes. Therefore, in this study, we attempted to examine the role of cholesterol and tension on the binding of PACSIN2 to membrane. We investigated the effect of cholesterol on the binding affinity of PACSIN2 F-BAR and on membrane shaping of liposomes composed of phospholipids with the specific fatty acids that are abundant in caveolae, which are POPC and POPS, in the presence or absence of membrane tension. We found that cholesterol reduced the affinity of PACSIN2 F-BAR to the liposomes, inhibiting the formation of straight membrane tubules. Consistently, the depletion of cholesterol from the plasma membrane resulted in enhanced tubule formation of PACSIN2 localization with caveolin-1, which appeared to represent intermediates of the removal of caveolae from the plasma membrane through caveolar endocytosis. The PACSIN2 knockout cells demonstrated that caveolar removal from the plasma membrane was dependent on PACSIN2. Therefore, this study indicates a novel role of PACSIN2 in the removal of cholesterol-less caveolae from the plasma membrane, which may represent an important mechanism for the maintenance of cholesterol-enriched caveolae on the plasma membrane.

## RESULTS

### The binding of PACSIN2 to the liposomes with or without cholesterol

The acute depletion of cholesterol can be tested by the application of cells with methyl-β-cyclodextrin (MβCD), which implies the requirement of two kinds of membranes – one with and one without cholesterol. To mimic the caveolar lipid membrane, we prepared liposomes using POPC, POPS, and cholesterol at a 33, 22, and 45 molar ratios, or by using POPC and POPS in 60 and 40 molar ratios, as discussed above. Membrane packing defects and charge density appeared to affect the binding of PACSIN2. To precisely describe the packing defects and charge density, we estimated the packing defects and charge density of the reconstituted membranes using molecular dynamics simulations (Figure 1A). We estimated the charge densities and the area of the packing defects of >10 Å^2^ within the simulated membrane. We also calculated the area of the membrane per lipid weight to compare the affinity of PACSIN2 per membrane area. The calculation indicated that the addition of cholesterol decreased the charge density while increasing the packing defects (Figure 1A).

**Figure 1.**
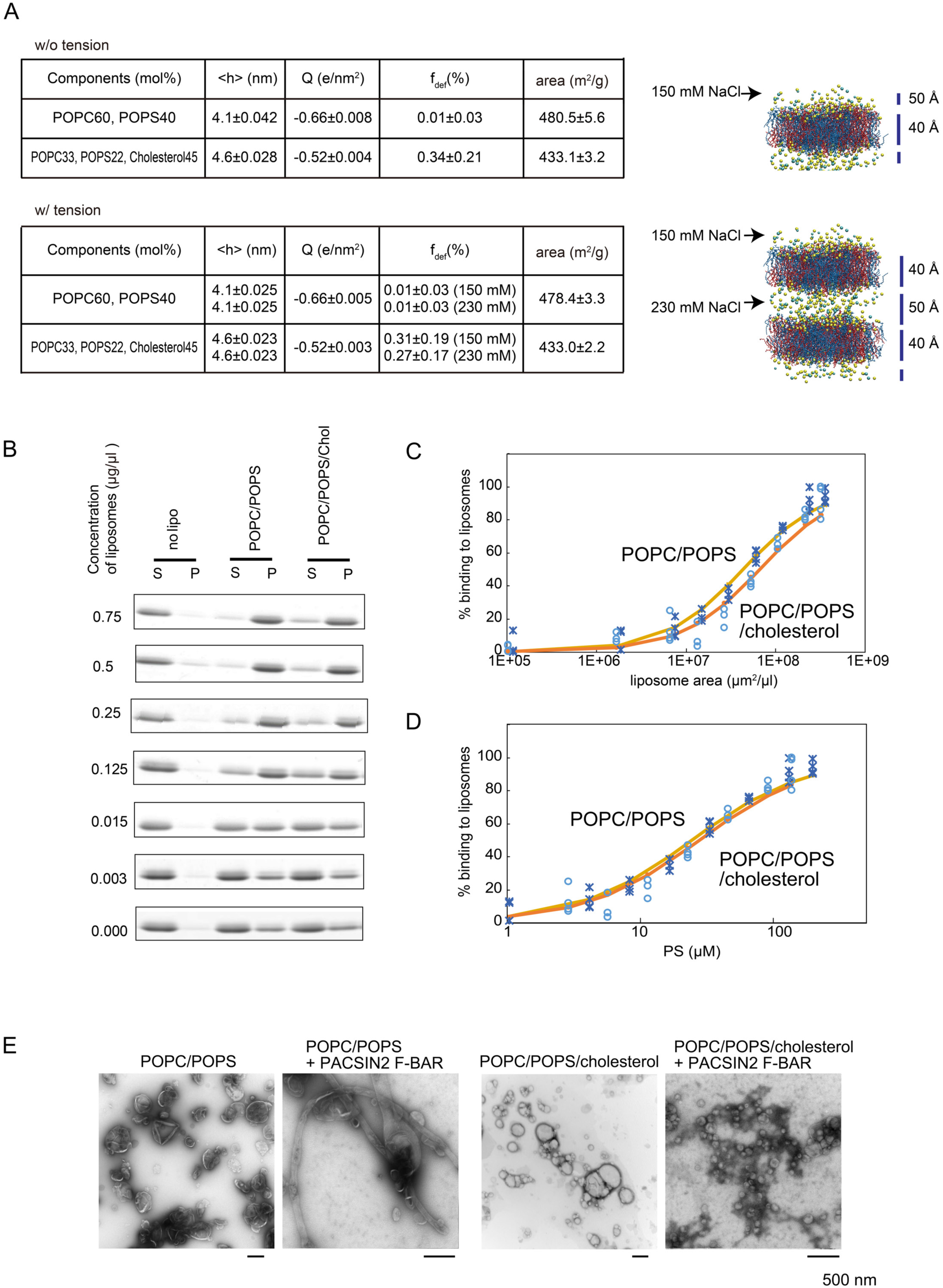
Effect of cholesterol on membrane binding of PACSIN2 F-BAR domain. (A) Molecular dynamics simulation for the estimation of model membrane parameters. The charge density (electron/nm^2^) was plotted against the average percentage of packing defects of >10 Å^2^ to the total membrane area (f_def_). (B-D) The binding of purified PACSIN2 F-BAR (5 µM) to liposomes comprised of POPC/POPS and POPC/POPS/cholesterol, determined by liposome co-sedimentation assay. PACSIN2 F-BAR was incubated with liposomes and the supernatant (S) was separated from the pellet (P) by centrifugation. S and P were subjected to SDS gel and stained using CBB. The binding graph represents the % binding of PACSIN2 F-BAR against the concentration of liposomes. Experiments were performed at least three times at each liposome concentration. The half maximum concentration of the binding was analyzed using Excel solver. (E) Electron microscopic images of negatively stained liposomes. The liposomes of POPC/POPS or POPC/POPS/cholesterol in the presence of PACSIN2 F-BAR domain (5 µM). Liposome concentration, 0.125 µg/µl.

The effect of cholesterol on the binding affinity of PACSIN2 F-BAR was investigated by determining the concentration of liposomes required for 50% binding using a liposome co-sedimentation assay (Figure 1B). From the amount of the PACSIN2 proteins in the liposomal pellet, we estimated the fraction of PACSIN2 that bound to the membrane. The percentages of PACSIN2 F-BAR binding to the liposomes in the co-sedimentation assay were plotted as a function of the membrane area and the molar concentration of PS (Figure 1C, D). All the membrane were assumed to be unilamellar for the plots. The PACSIN2 F-BAR domain binding to the POPC/POPS liposomes reached up to 50% when the total membrane area was 4.3 × 10^7^ µm^2^/µl, while binding to POPC/POPS/cholesterol liposomes reached 50% when the total membrane area of was 6.5 × 10^7^ µm^2^/µl, suggesting that the presence of cholesterol decreased the affinity of the PACSIN2 F-BAR domain to the membrane (Figure 1C). Interestingly, when we examined the 50% binding with respect to the DOPS concentration, the binding of the PACSIN2 F-BAR domain to the POPC/POPS liposomes reached 50% when the DOPS concentration was 23 µM in the absence of cholesterol, and at 27 µM in the presence of cholesterol (Figure 1D). The difference in the concentration for 50% binding was smaller as a function of the PS concentration compared to that of the area. This could be explained by the decrease in the charge density (Figure 1A). On the other hand, the increase in the packing defect upon the addition of cholesterol might not contribute to the affinity of PACSIN2 to the membrane; rather, the charge density in the presence of cholesterol may account for the difference in the membrane binding of PACSIN2 in a cholesterol-dependent manner.

### Membrane deformation by PACSIN2 is cholesterol-dependent

In order to investigate whether cholesterol influenced the membrane remodeling of PACSIN2, we analyzed the morphology of liposomes in the presence of PACSIN2 F-BAR using transmission electron microscope (TEM). The POPC/POPS liposomes were found to be deformed by PACSIN2 F-BAR into straight tubular shapes (Figure 1E). However, the POPC/POPS/cholesterol liposomes were remodeled into tubules resembling an assembly of small vesicles (Figure 1E), similar to “beads on a string”, which correlates with previous observations in bovine Folch liposomes (Wang et al., 2009). These results indicated that cholesterol suppressed the formation of straight tubules by PACSIN2 F-BAR, which might play a role in the formation of the caveolar shape.

### Non-altered membrane binding of PACSIN2 upon hypotonic tension *in vitro*

In the previous study, PACSIN2 was found to respond to membrane tension that was applied by hypotonic osmolarity (Senju et al., 2015). PACSIN2 was found to be removed from the caveolae upon hypo-osmotic treatment of the cells, presumably due to a phosphorylation-induced weakening of the membrane binding as a result of the introduction of a negative charge. However, it is possible that the tension, tension-induced packing defects, and tension-induced resistance to deformation may modulate the binding of PACSIN2 to the cell membranes. Here, using hypo-osmotic tension, we examined the effect of tension on the binding and remodeling activity of PACSIN2 F-BAR to liposomes. Osmotic pressure induces the stretching of liposomes, which eventually causes liposomes to swell and rupture (Alam Shibly et al., 2016; Finkelstein et al., 1986). We determined the tension threshold at which liposomes could exist without rupturing by observing the leakage of the fluorescent molecules from the liposomes. The liposomes were constructed using buffer containing carboxyfluorescein (CF), and then purified using gel filtration and centrifugation to remove any CF found outside the liposomes. The liposomes were then treated with a hypotonic buffer to induce membrane tension. High hypotonic tension can lead to liposome rupture, followed by leakage of the internal CF, which can be quantified by the spectro-fluorometry analysis of CF released from the liposomes (Hamai et al., 2007; Shoemaker and Vanderlick, 2002). Here, we examined the release of CF from the liposomes made of porcine brain Folch fraction, a total fraction of porcine brain, under several levels of hypotonic tension (Figure 2A). The CF fluorescence was found to increase as the osmolarity difference increased, suggesting that the rupture of the liposomes increased as the tension increased. Therefore, we selected 435 mOsmol sucrose as the solute inside liposomes, and 300 mOsmol glucose buffer as the external solute under the same salt concentrations to obtain an osmolarity difference of 135 mOsmol. Then, we applied this tension to the POPC/POPS liposomes and to POPC/POPS/cholesterol liposomes and observed the leakage. The leakage was considered to be similar to that of porcine Folch liposomes (Figure 2B).

**Figure 2.**
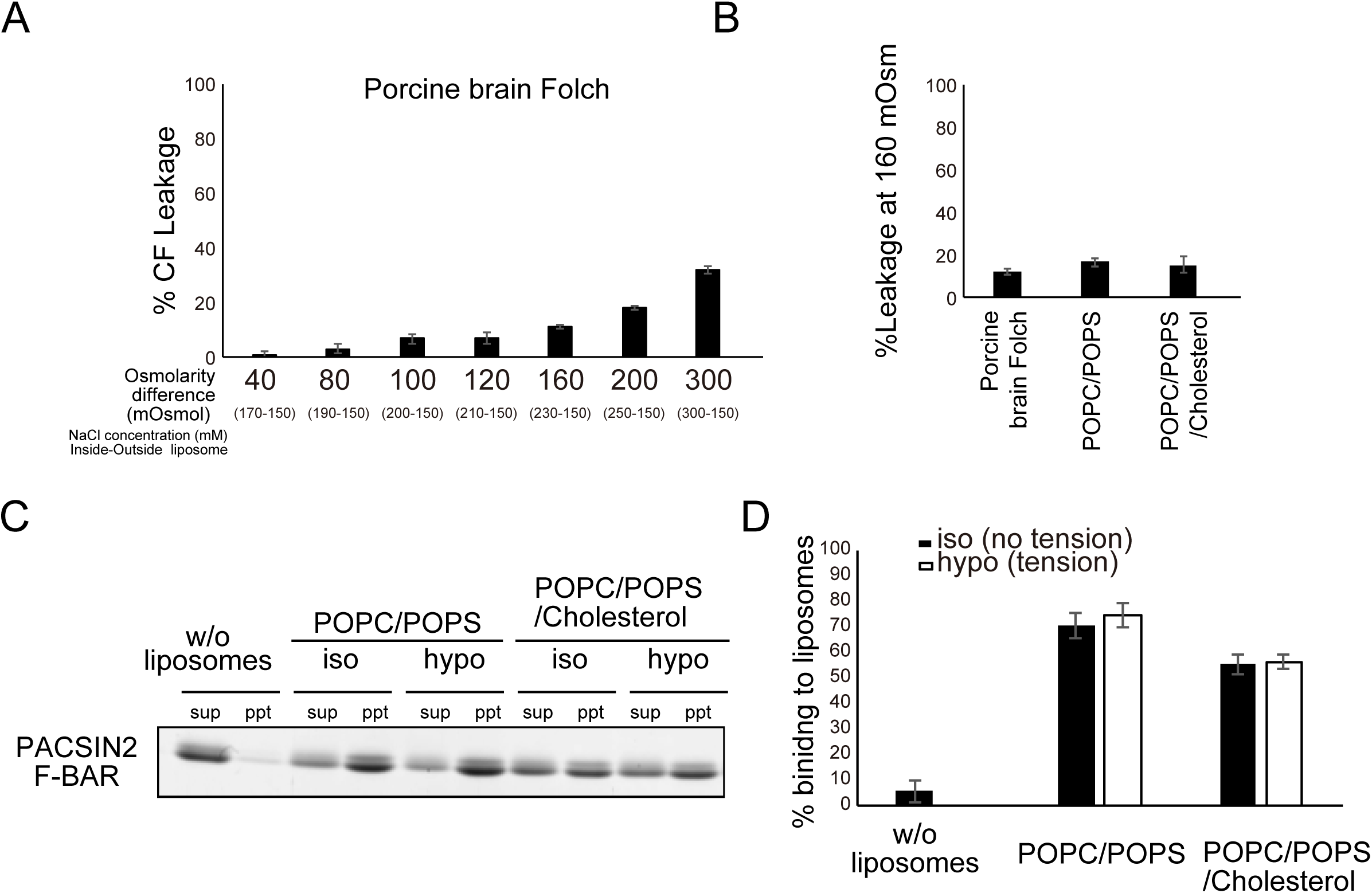
Osmolality and membrane binding of PACSIN2 F-BAR domain. (A) % of carboxyfluorescein (CF) released from porcine brain Folch liposomes upon exposure to different levels of tension. (B) % of CF released from several types of liposomes at a tension induced by 160 mOsmol difference between the inside and outside of the liposomes. Experiments were performed three times. Error bar indicates the SD. (C) Binding affinity of PACSIN2 F-BAR (5 µM) to POPC/POPS and POPC/POPS/cholesterol under isotonic (no tension) and hypotonic tension by co-sedimentation assay. PACSIN2 F-BAR was incubated with liposomes and the supernatant (S) was separated from the pellet (P) by centrifugation. S and P were subjected to SDS gel and CBB staining. Liposome concentration, 0.125 µg/µl. (D) Quantification of (C). The assay was performed 4 times. Error bar indicates the SD. Statistically significance between the isotonic and hypotonic condition was analyzed using paired Student’s *t*-test.

The packing defects upon tension application was also examined by molecular dynamic simulations. The number of ions were adjusted according to the experimental setup above to mimic the difference in osmolarity. Then the changes in the packing defects were not observed (Figure 1A). Therefore, the tension without rupture could not induce the packing defect, and the formation of packing defect by membrane tension was considered to be negligible.

We then examined the binding affinity of PACSIN2 F-BAR to POPC/POPS and POPC/POPS/cholesterol liposomes under both isotonic and hypotonic tension using a liposome co-sedimentation assay (Figure 2C). Since the amount of the PACSIN2 proteins in the liposomal pellet was similar independent of tension, the binding affinity of PACSIN2 F-BAR under both isotonic and hypotonic tension appeared to be similar (Figure 2D), suggesting that the membrane tension below the rupture induction could not alter the amount of PACSIN2 F-BAR domain on the liposomes.

### Acute depletion of cholesterol induced the tubulation of PACSIN2 in HeLa cells

We then examined whether the amount of cholesterol could modulate the membrane tubules of PACSIN2 localization in cells. Since the overexpression of the PACSIN2 F-BAR domain induced massive tubulation in the cells (Senju et al., 2011), the HeLa cells were weakly expressed with GFP-PACSIN2 F-BAR domain, and then observed using a microscope during the addition of MβCD to deplete cholesterol from the plasma membrane. The depletion of cholesterol induced the acute tubule formation of PACSIN2 F-BAR domain localization (Figure 3A), which appears to be the consistent with the observations of the *in vitro* tubulation of PACSIN2.

**Figure 3.**
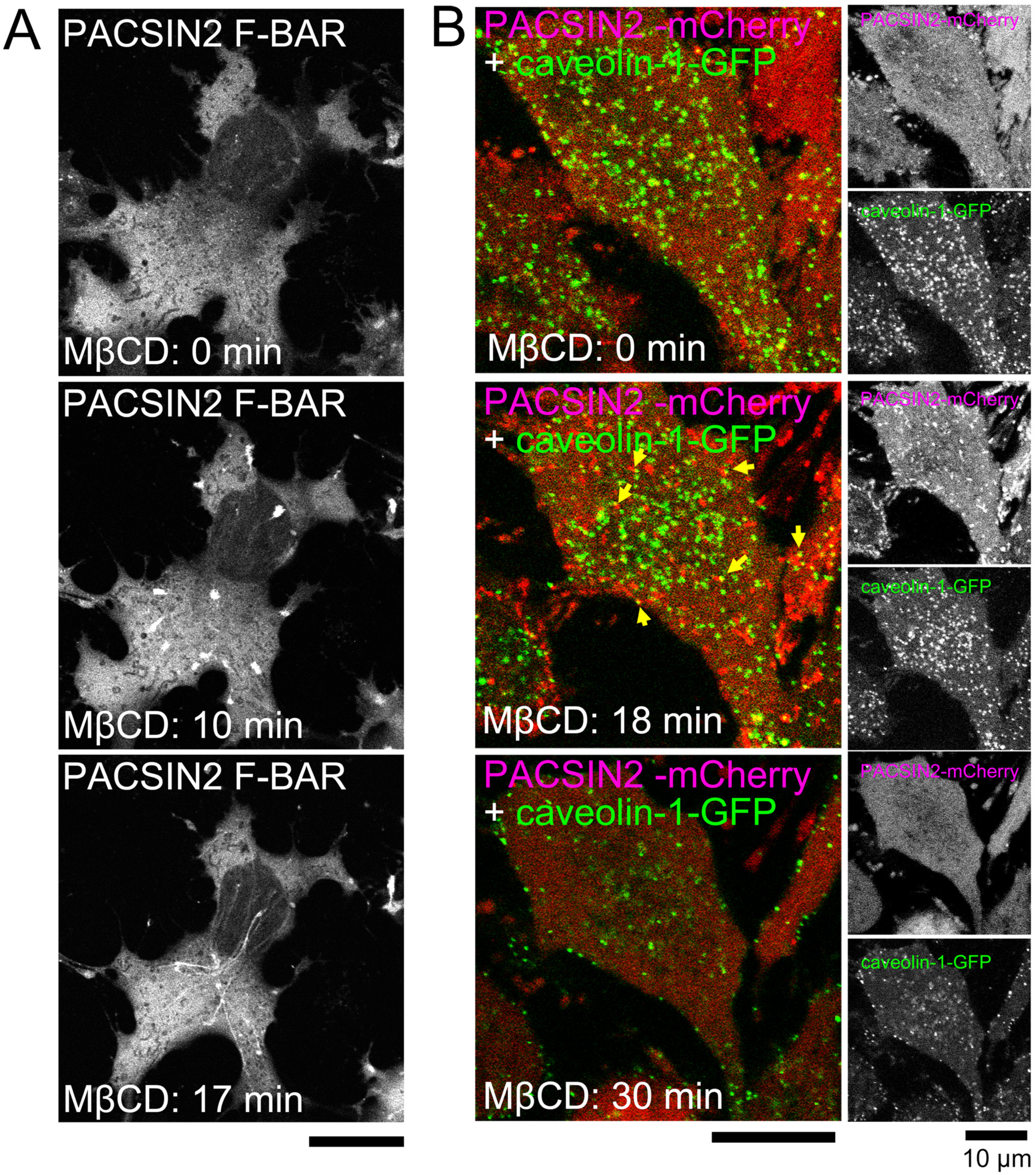
Formation of PACSIN2 tubules upon cholesterol depletion. (A) HeLa cells expressing PACSIN2 F-BAR domain fragment showing acute tabulations after treatment with methyl-β-cyclodextrin (MβCD). The tubulation was observed to appear at 13 min after the cells were treated with MβCD. (B) HeLa cells expressing PACSIN2-mcherry and caveoln-1-GFP after (MβCD) treatment show acute tubulation with caveolin-1 at the tip of the tubules. PACSIN2 were transiently recruited to the caveolin-1 associated membrane (shown in the blue box, denoted by white arrows) and impaired the morphology of caveolae, followed by a decrease in the number of caveolae.

Next, we examined the behavior of full-length PACSIN2 and caveolin-1 upon treatment with MβCD. Caveolin-1-GFP and PACSIN2-mCherry were expressed at endogenous protein levels. Upon MβCD treatment, acute tubules of PACSIN2-mCherry were observed, with the tips of the tubules sometimes staining positive for caveolin-1-GFP (Figure 3B). These findings suggested that caveolar invagination was elongated into tubules upon cholesterol depletion by PACSIN2. Interestingly, the tubules and caveolin-1 eventually disappeared, suggesting that the caveolin-1 proteins or caveolae were internalized upon cholesterol depletion by MβCD. Therefore, PACSIN2 was suggested to mediate caveolar internalization upon cholesterol depletion via its tubule-forming ability.

### PACSIN2-mediated internalization of caveolin-1 upon cholesterol depletion

The amount of caveolae in cells has been shown to be dependent on cholesterol (Hailstones et al., 1998). However, the mechanism for cholesterol-dependent caveolar regulation is unknown. We prepared PACSIN2 knockout cells using CRISPR/Cas9 and determine the levels of caveolin-1 upon depletion of cholesterol by MβCD. The cells were seeded in serum-free medium and treated with MβCD for 3 h. After MβCD treatment, the amount of caveolin-1 relative to GAPDH was found to be lower in the PACSIN2 knockout cells compared with that in the parental HeLa cells, suggesting that the regulation of the amount of caveolin-1 is dependent on PACSIN2 (Figure 4A, B).

**Figure 4.**
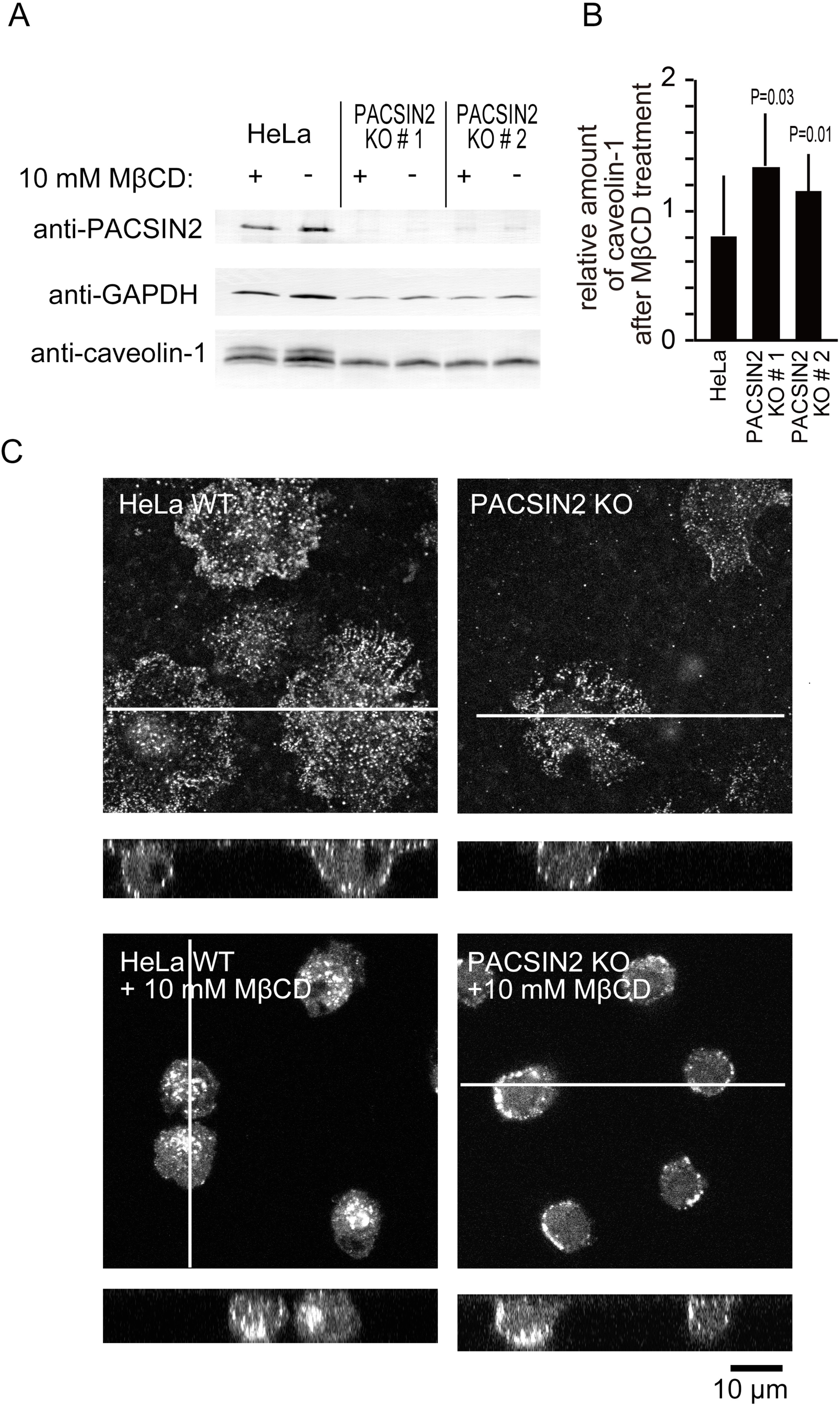
Internalization of caveolin-1 upon depletion of cholesterol by PACSIN2. (A) HeLa cells and the two lines of PACSIN2 knockout HeLa cells were treated with 10 mM MβCD for 3 hr. The amounts of PACSIN2, caveolin-1, and GAPDH were analyzed using Western blotting. (B) Quantification of (A), showing alterations of the amount of caveolin-1 upon MβCD treatment. The amount of caveolin-1 relative to GAPDH was examined and the fold changes by MβCD are shown. (C) Caveolin-1 distribution in HeLa cells and PACSIN2 knockout HeLa cells after treatment with MβCD (10 mM). The cells were fixed and labelled with anti-caveolin-1 antibody. The white line indicates the sectioning plane in the lower panels.

The localization of caveolin-1 in both HeLa cells and the PACSIN2 knockout cells was then examined by immuno-staining using an antibody against caveolin-1. The localization of caveolin-1 in PACSIN2-knockout cells was similar to that of parental HeLa cells under confocal microscope. After 3 h of MβCD treatment, the cellular localization of caveolin-1 was examined. Caveolin-1 was internalized from the plasma membrane in HeLa cells (Figure 4C). Importantly, caveolin-1 remained on the plasma membrane in the PACSIN2 knockout cells after MβCD treatment (Figure 4C), suggesting that PACSIN2 mediated the internalization of the caveolin-1 from the plasma membrane upon cholesterol depletion. These results suggest that PACSIN2 mediates the internalization of caveolin-1 in the plasma membrane upon the depletion of cholesterol.

## DISCUSSION

In this study, we found that the membrane binding of PACSIN2 was weaker when the membrane contained cholesterol, using the reconstituted membrane containing abundant lipids in caveolae such as palmitoyl-oleoyl (PO) PC, POPS, and cholesterol. This observation suggested that cholesterol depletion strengthened the membrane binding of PACSIN2, thereby inducing tubule formation of caveolae and resulting in the internalization of caveolae, presumably through their endocytosis, i.e., the scission of the tubules (Figure 5).

**Figure 5.**
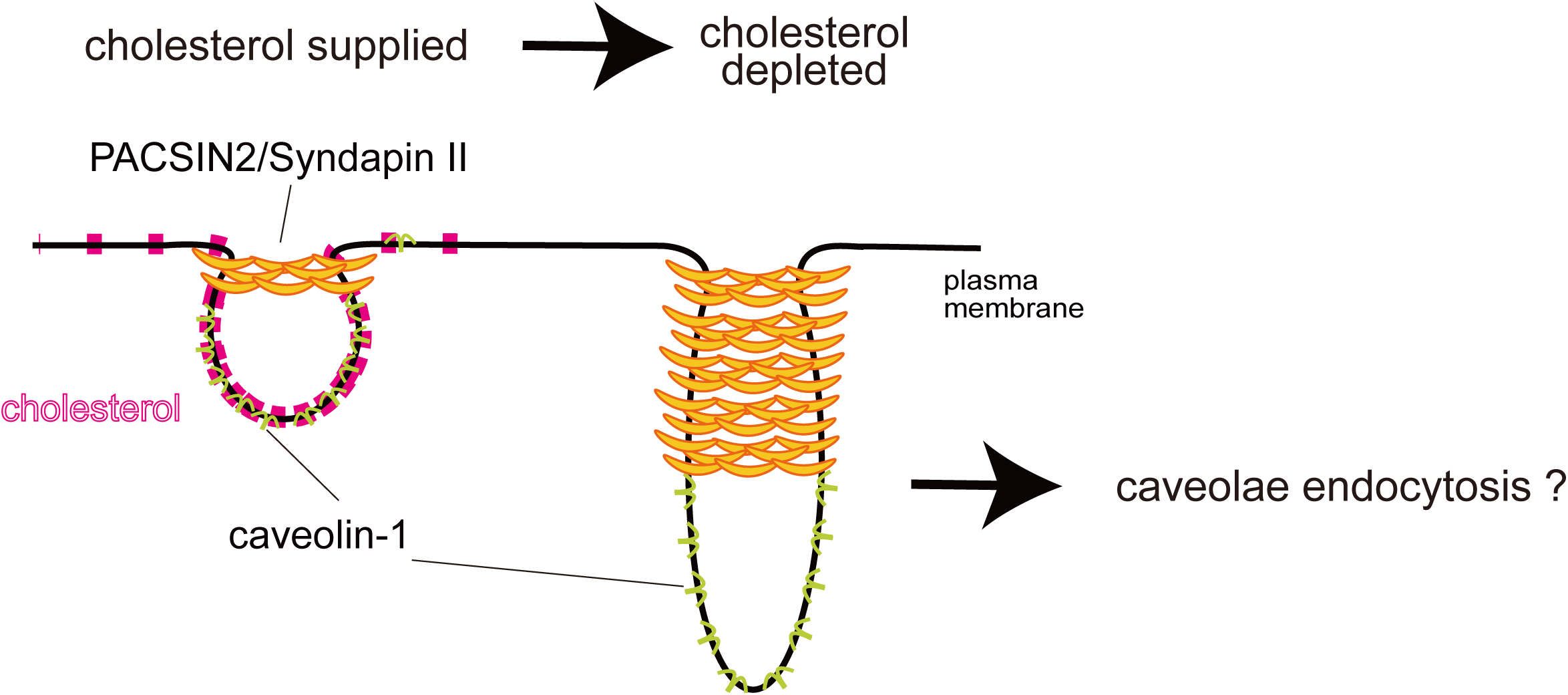
Potential model of PACSIN2-induced tubules for caveolar morphology. Membrane tubulation by PACSIN2 was enhanced in the cholesterol-less membrane. In contrast, membrane tubulation by PACSIN2 was not frequently observed in the cholesterol-containing membrane. Therefore, upon cholesterol depletion, PACSIN2-localized caveolae were tubulated, which may represent the intermediate of endocytosis. Upon cholesterol depletion, the caveolae were internalized in a PACSIN2-dependent manner.

The localization of PACSIN2 at the neck of caveolar invaginations, but not at the entire caveolar flask (Senju et al., 2011), appeared to be consistent with the negative regulation of membrane binding by cholesterol, because the neck region at the boundary of caveolae was supposed to have smaller amounts of cholesterol than the caveolar bulb. Interestingly, our electron microscopic observations of liposomes with or without cholesterol in the presence of PACSIN2 indicated that cholesterol could modulate the membrane shaping ability of PACSIN2. The POPC/POPS/cholesterol liposomes were deformed by the PACSIN2 F-BAR domain into a “beads on a string” morphology, which resembles the curvature of the flask-shaped caveolae. This “beads on a string” morphology was previously reported in bovine brain Folch liposomes, which contain cholesterol (Wang et al., 2009). In contrast, the POPC/POPS liposomes without cholesterol were deformed by the PACSIN2 F-BAR domain into straight tubules, which was indicative of tubule formation in the cells. It is also possible that the cholesterol in the membrane changes the deformability of the membrane, as indicated by the differential amounts of the packing defect (Figure 1A). However, differences in the affinity of the PACSIN2 F-BAR domain to the membrane may also account for the differential morphology.

The differences in binding appeared to be correlated to the presence of cholesterol in the membrane. Molecular dynamics simulation suggested that the presence of cholesterol can increase the packing defects in the membrane, while tightening the packing of the entire membrane, which is indicated by the tight packing of phospholipids by the association of cholesterol with the acyl-chain of phospholipids (Falck et al., 2004; Lund-Katz et al., 1988; Presti et al., 1982). Due to the presence of the hydrophobic loops thought to be inserted into the packing defects of membrane (Shimada et al., 2010), the increase in packing defects was first hypothesized to lead to an increase in the binding of PACSIN2 to the cell membrane. The increase of packing defects may occur upon the application of tension to the membrane. However, the results of molecular dynamics simulation and leakage assay indicated that the tension before membrane rupture did not increase the packing defect nor the rate of PACSIN2 binding to the membrane. Although the mutation at the loop resulted in reduced binding of PACSIN2 to the membrane (Shimada et al., 2010), our results suggested that the charge density could be the primary determinant of membrane binding. The decreased affinity of PACSIN2 to the membrane by the depletion of cholesterol may be explained by the decrease in the charge density resulting from the addition of cholesterol.

It is widely known that caveolae have a cholesterol-dependent structure (Fielding and Fielding, 2000; Harder and Simons, 1997; Hooper, 1999). However, the mechanisms of the cholesterol-dependent regulation of caveolae have not yet been elucidated. In this study, the formation of tubules as a result of cholesterol depletion suggest a mechanism of caveolar downregulation upon cholesterol depletion.

## MATERIALS AND METHODS

### Molecular dynamics

Initial bilayers containing 400 lipids/cholesterol molecules (200 molecules in each leaflet), water molecules with a thickness of 22.5 Å at the top and bottom of the bilayers, and 150 mM NaCl were constructed using a CHARMM-GUI membrane builder (Wu et al., 2014). The lipid/cholesterol compositions of the simulated systems are shown in Figure 1A. We also constructed two bilayer systems (with tension in Figure 1A) to investigate the effects of tension induced by the osmotic pressure. After 500-ns MD simulations, the single bilayer systems were replicated along the bilayer normal direction. Then, randomly selected water molecules were replaced with Na^+^ or Cl ^−^ so that the two regions divided by two bilayers had different concentrations of NaCl (150 and 230 mM). All MD simulations were performed using Gromacs 2018.1 (Abraham et al., 2015; Berendsen et al., 1995). The system was brought to thermodynamic equilibrium at 300 K and 1 atm using the Nosé-Hoover thermostat and the Parrinello-Rahman barostat with semi-isotropic pressure control. The equations of motion were integrated with a time step of 2 fs. Long-range Coulomb energy was evaluated using the particle mesh Ewald (PME) method. The CHARMM36m force field (Huang et al., 2017) was used. Simulations were conducted for 500 ns, and trajectories were saved every 0.1 ns.

To detect the packing defects, we followed the procedure by Vamparys et al. (Vamparys et al., 2013). We used 4 Å for vdW radii of carbon and phosphorus atoms and 3 Å for those of oxygen and nitrogen atoms. The last 300-ns (3,000 snapshots) were used for the analysis. To evaluate a fraction of the defect area, we defined a fraction as the sum of the defect area equal to or larger than 10 Å^2^.

### GST-tagged protein purification

Rosetta Gami B-containing pGEX6P1-PACSIN2 constructs (Shimada et al., 2010) were cultured in LB medium overnight at 20°C after the addition of IPTG. GST-tagged expressed proteins were purified using GST beads (Glutathione Sepharose 4B; GE Healthcare Life Science). On the following day, the cells were centrifuged and the resulting pellets were stored at ‒80°C as the stocks. Before purification, GST beads were washed with *E. coli* sonication buffer (10 mM Tris HCl pH 7.5, 150 mM NaCl, 1 mM EDTA, 0.5% Triton X-100, Milli Q water) three times. To purify the proteins, the cell pellets were resuspended in the mixture of the sonication buffer, 0.1% DTT, and 1% PMSF, followed by sonication in an ice container to disrupt the cell membrane (3-s burst – 3-s rest for total time 4 min). The lysates were then centrifuged at 15,000 rpm for 10 min at 4°C. The supernatant was transferred into a suspension of GST beads and rotated for 1 h at 4°C. The beads were then centrifuged at 500 *g* for 5 min at 4°C, the supernatant was discarded, and the beads were suspended in the wash buffer (10 mM Tris HCl pH 7.5, 150 mM NaCl, 1 mM EDTA). The washing of the beads was repeated over three times. Subsequently, PreScission Protease (GE Healthcare) was added to remove GST from the proteins, followed by overnight rotation at 4°C. The protein was then separated from the GST beads using a mini column before storage as a stock solution at ‒80°C. The purified proteins were visualized using an SDS-PAGE gel and Coomassie Brilliant Blue (CBB) staining. The protein concentration was determined using the band intensities of the gels in Image J.

### Liposome preparation

Liposomes were prepared from 1-palmitoyl-2-oleoyl-sn-glycero-3-phosphocholine (POPC), 1-palmitoyl-2-oleoyl-sn-glycero-3-phospho-L-serine (POPS), and cholesterol. These lipids were purchased from Avanti polar lipids. To obtain the liposomes, the lipids were dried under nitrogen gas, followed by drying in a vacuum for at least 1 h to remove any traces of residual solvent. Then, 435 mOsmol/kg sucrose buffer (435 mM sucrose, 10 mM Tris HCl pH 7.5, 1 mM EDTA) was added to the thin layer of dried lipids, followed by vortexing to induce liposome formation. The liposome suspension was subjected to ten freeze-thaw cycles in liquid nitrogen, followed by thawing in water bath at 45°C and either storage at ‒30°C or immediate use in the assays. Before use, the liposomes were extruded through the polycarbonate membrane with a pore size of 2 µm.

### Liposome co-sedimentation assay

The liposomes used in this assay were composed of POPC/POPS and POPC/POPS/cholesterol. Proteins were mixed with the liposomes. The osmolarity outside the liposomes was adjusted using the buffer of various concentrations of glucose (0-135 mM glucose, 10 mM Tris HCl pH 7.5, 1 mM EDTA, 150 mM NaCl) to obtain isotonic (435 mOsmol/kg inside and outside liposome) and hypotonic conditions (435 mOsmol/kg inside and 300 mOsmol/kg outside liposome). The mixture was then incubated at room temperature for 20 min, followed by centrifugation at 50,000 rpm for 20 min at 25°C in a TLA100 rotor (Beckman Coulter). Following these procedures, any bound proteins would be found in the pellet, while unbound proteins would be found in the supernatant. The pellet and supernatant were then separated, subjected to SDS PAGE, and stained using CBB.

### Transmission electron microscopy (TEM)

The proteins and liposomes were prepared and mixed as in the liposome co-sedimentation assay, then placed on a parafilm surface on a heating block at 25°C. After incubation for 20 min, the samples were placed on a grid (Nisshin EM) covered with Polyvinyl Formal (Formvar; Nisshin EM). NaCl was then removed using HEPES buffer, and the samples were stained with 0.5% uranyl acetate before air-drying for several hours to remove the remaining solutions. Dried sections were then observed using a transmission electron microscope (Hitachi H-7100).

### Osmotically induced leakage assay

The lipids used in these experiments were porcine brain Folch, 1-palmitoyl-2-oleoyl-sn-glycero-3-phosphocholine (POPC), 1-palmitoyl-2-oleoyl-sn-glycero-3-phospho-L-serine (POPS), and cholesterol dissolved in chloroform (Avanti Polar Lipids). These lipids were dried under nitrogen gas, followed by drying in vacuum for 20 min. The dried lipids were suspended in 300 µl of the buffer containing the indicated concentrations of NaCl, 10 mM Tris HCl pH 7.5, 1 mM EDTA, and 20 µM 5(6)-carboxyfluorescein (CF) for 1-2 h. The osmolarity inside and outside the liposomes was adjusted using the concentration of NaCl. Alternatively, the dried lipids were solubilized in the buffer containing 460 mM sucrose, 10 mM Tris HCl pH 7.5, 1 mM EDTA, and 40 µM CF. Then, 10 mL of Sephadex G-50 was suspended in the buffer that was used to prepare liposomes but without CF. The Sephadex G-50 suspension was then settled in a chromatography column. The liposomes were added to the column, and the CF outside the liposomes was removed. The eluate drops were collected in Eppendorf tubes until the elution of the CF phase. The tubes containing liposomes loaded with CF were verified under a UV light. These liposomes were then collected in a single Eppendorf tube and divided into three treatment groups: isotonic conditions, hypotonic conditions, and liposomes for the determination of total CF. All of the groups were centrifuged at 50,000 rpm for 20 min, and the resulting supernatant was removed. The pellet of the first group was resuspended with isotonic NaCl or glucose buffer, while the pellet of the second group was resuspended with hypotonic buffer. Both groups were incubated at room temperature for 20 min, followed by centrifugation at 50,000 rpm for 20 min. The fluorescence intensity of the supernatant was measured using a spectrofluorometer (Jasco FP-6500). The third group was resuspended using the buffer with detergent (10 mM Tris HCl pH 7.5, 150 mM NaCl, 1 mM EDTA, 0.5% Triton X-100) for at least 10 min to allow for the release of CF from the liposomes. The fluorescence intensity of CF was measured as the percentage of the total CF. Total CF leakage from liposome was calculated using the following formula:

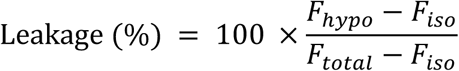

where F_*hypo*_, F_*iso*_, and F_*total*_ denote the CF intensity of hypotonic, isotonic, and total CF in liposomes, respectively. The buffer osmolality was confirmed using a Wescor Vapro 5600 vapor pressure osmometer, according to manufacturer’s protocol. The unit of measurement was mOsmol/kg.

### Cell culture and MβCD treatment

HeLa cells were cultured in Dulbecco’s modified Eagle medium (DMEM) (Nacalai) supplemented with 10% fetal calf serum (FCS), penicillin, and streptomycin (Meiji pharmaceuticals). For MβCD treatment, the HeLa cells were suspended in DMEM supplemented with 1/100 concentration of insulin, transferrin, and selenium solution (ITS-G) (Gibco 41400045) and cultured overnight. MβCD (Sigma) was then added to the cells at 10 mM for 3 h.

### Expression of PACSIN2 in HeLa cells

EGFP-labeled PACSIN2 in pEGFP-C1 (Senju et al., 2015) was transfected using Lipofectamine 3000 and PLUS reagents (Invitrogen), according to the manufacturer’s instructions. mCherry-labeled PACSIN2 was prepared by subcloning PACSIN2 cDNA into pmCherry-C1 vector (Clontech), in which the GFP in pEGFP-C1 was replaced with mCherry. Cells stably expressing caveolin-1-EGFP were prepared as previously described (Senju et al., 2015). The stable expression of PACSIN2-mCherry into the caveolin-1-EGFP expressing HeLa cells (Moren et al., 2012) was performed using the pMXs vector and retrovirus that was produced in the packaging cell line Plat-A (Kitamura et al., 2003). After the FACS sorting and cloning of the cells, the cells with PACSIN2-mCherry expression equivalent to the endogenous PACSIN2 level were selected for observation using western blotting.

### Live observation

For live observation using total internal reflection microscopy, HeLa cells were grown on a glass-bottomed dish using DMEM containing 10% FBS and 10 mM HEPES (pH 7.5). Images were acquired for 5 min at 1 or 0.5 s intervals using a confocal microscope (Olympus FV1000D) equipped with a 100× NA 1.45 oil immersion objective (Olympus). Live cells were treated with 10 mM methyl-β-cyclodextrin (MβCD).

### PACSIN2 knockout cells

The guide RNA targeting the first exon of PACSIN2 (TGAGCGGGCGCGCATCGAGA) was designed using the MIT CRISPR server (http://crispr.mit.edu) (Hsu et al., 2013) and inserted into the pX459 vector (Ran et al., 2013) After transfection into HeLa cells, the cells were cloned by puromycin resistance.

### Immunofluorescent staining

Cells grown on poly-L-lysin-coated cover glasses were fixed with 4% paraformaldehyde with 0.2% glutaraldehyde in HEPES-buffered saline, containing 30 mM HEPES (pH 7.5), 100 mM NaCl, and 2 mM CaCl_2_ for 5 min. The cells were then blocked with DMEM supplemented with 10% fetal calf serum (FCS) overnight and incubated with anti-caveolin-1 antibody (BD610406) in 1% BSA in TBS (100 mM Tris-HCl (pH 7.5), and 150 mM NaCl) for 1 h. After washing with PBS, the cells were incubated with fluorescently labeled secondary antibody. The cells were washed again before mounting using Prolong Gold (Thermo Fisher/Invitrogen). Cell images were obtained using an FV1000 laser-scanning confocal microscope (Olympus) equipped with a 100× NA 1.45 oil lens (Olympus) at room temperature.

### Statistical analysis

All experiments were performed at least in triplicate. The results were presented as the mean ± standard deviation (SD). Significant data were analyzed using one-tailed (isotonic and hypotonic results) or two-tailed Student’s t-test (with and without cholesterol). The half maximum of the binding was determined using Excel solver. P<0.05 was considered statistically significant.

## ACKNOWLEDGMENT

This work was supported by grants from the Funding Program for Next Generation World-Leading Researchers (NEXT program LS031), JSPS (KAKENHI 26291037, JP15H0164, JP15H05902, JP17H03674, JP17H06006), JST CREST (JPMJCR1863), and the NAIST Interdisciplinary Frontier Research Project to S.S. This research was supported by MEXT/JSPS KAKENHI (JP19H03191) and MEXT Priority Issues on Post-K Computer Projects “Building Innovative Drug Discovery Infrastructure through Functional Control of Biomolecular System” to A.K. The computations were partly performed using the supercomputers at the RCCS at the National Institute of Natural Science and ISSP at the University of Tokyo. This research also used computational resources of the K computer provided by the RIKEN Advanced Institute for Computational Science through the HPCI System Research project (Project ID: hp190181).

